# ^13^C and ^15^N Resonance Assignments of Alpha Synuclein Fibrils Amplified from Lewy Body Dementia Tissue

**DOI:** 10.1101/2023.01.09.523339

**Authors:** Alexander M. Barclay, Dhruva D. Dhavale, Collin G. Borcik, Moses H. Milchberg, Paul T. Kotzbauer, Chad M. Rienstra

## Abstract

Fibrils of the protein α-synuclein (Asyn) are implicated in the pathogenesis of Parkinson Disease, Lewy Body Dementia, and Multiple System Atrophy. Numerous forms of Asyn fibrils have been studied by solid-state NMR and resonance assignments have been reported. Here, we report a new set of ^13^C, ^15^N assignments that are unique to fibrils obtained by amplification from postmortem brain tissue of a patient diagnosed with Lewy Body Dementia.

## Biological Context

Alpha-synuclein (Asyn) is a 140-residue protein found in the presynaptic termini of neurons in the brain (Clayton and George 1999). While elucidation of the exact function of this protein in the brain remains elusive (Lautenschlager et al. 2018), the aggregation of Asyn in the form of fibrils has been a pathological hallmark of Parkinson Disease (PD), Lewy Body Dementia (LBD) and Multiple System Atrophy (MSA), all which can be classified as α-synucleinopathies.

Dementia occurs frequently in PD, sometimes beginning at approximately the same time as motor symptoms (Dementia with Lewy bodies or DLB), or up to 20 years after motor symptoms begin (PD with dementia or PDD). The term LBD encompasses the spectrum of clinical presentations classified as DLB and PDD.

Asyn fibrils have been reported to have a range of tertiary and quaternary structures (Schweighauser et al. 2020, Tuttle et al. 2016), contributing to an emerging understanding of the precise relationships of *in vitro* and *in vivo* conditions. The *in vitro* structures depend on various factors such as mutations (Comellas et al. 2011, Khalaf et al. 2014, Kruger et al. 1998, Lemkau et al. 2012, Lemkau et al. 2013, Polymeropoulos et al. 1997, Zarranz et al. 2004), cytosolic components (Guilarte 2010, Kwakye et al. 2015, Shin and Chung 2012), lipids (Bodner et al. 2009, Bodner et al. 2010, Jakubec et al. 2021, Mahapatra et al. 2021), and metals (Uversky et al. 2001). Growing evidence indicates that distinct polymorphs are associated with pathologic Asyn accumulation in α-synucleinopathies, as detailed by previously reported *in vivo* studies (Frieg et al. 2022, Schweighauser et al. 2020, Yang et al. 2022). Therefore, structural determination of these fibrils is vital for advancing the understanding of disease etiology, and to aid the development of polymorph-specific clinical diagnostic tools and novel therapeutics.

We isolated insoluble Asyn fibrils from postmortem LBD tissue. Then, to analyze LBD fibril structure by SSNMR, we amplified the Asyn fibril seeds using uniform [^13^C, ^15^N] labeled wild-type Asyn. Here, we report the ^13^C and ^15^N chemical shifts for the amplified Asyn fibrils from an LBD autopsy case.. The spectra of these *ex vivo* fibrils exhibit resonances that differ from those of previously reported *in vitro* fibril preparations. These findings demonstrate a different arrangement of β-strands, supporting the hypothesis that fibril structure is directly linked to disease phenotype.

## Methods and experiments

### Protein expression and purification

Expression of uniform [^13^C, ^15^N] labeled wild-type Asyn was carried out in *E. coli* BL21(DE3)/pET28a-AS in modified Studier medium M (Studier 2005). The labeling medium contained 3.3 g/L [^13^C]glucose, 3 g/L [^15^N]ammonium chloride, 11 mL/L [^13^C, ^15^N]Bioexpress (Cambridge Isotope Laboratories, Inc., Tewksbury, MA), 1 mL/L BME vitamins (Sigma), and 90 μg/mL kanamycin. After a preliminary growth in medium containing natural abundance (NA) isotopes, the cells were transferred to the labeling medium at 37 ºC to an OD_600_ of 1.2, at which point the temperature was reduced to 25 ºC and protein expression induced with 0.5 mM isopropyl β-D-1-thiogalactopyranoside (IPTG) and grown for 15 h to a final OD_600_ of 4.1 and harvested.

Protein purification was done as described previously (Barclay et al. 2018). Briefly, cells were lysed chemically in the presence of Turbonuclease (Sigma) to digest nucleic acids. Purification began with a heat denaturation of the cleared lysate, followed by ammonium sulfate precipitation (Kloepper et al. 2006). The resolubilized protein was bound to QFF anion exchange resin (GE Healthcare Life Sciences, Marlborough, MA) and eluted using a linear gradient of 0.2–0.6 M NaCl. Fractions containing Asyn monomer, which eluted at about 0.3 M NaCl, were pooled, concentrated, and run over a 26/60 Sephacryl S-200 HR gel filtration column (GE Healthcare Life Sciences) equilibrated in 50 mM Tris-HCl, 100 mM NaCl, pH 8 buffer. Fractions were pooled, concentrated to ∼20 mg/mL Asyn, and dialyzed at 4 ºC into 10 mM Tris-HCl pH 7.6, 50 mM NaCl, 1 mM DTT, and stored at a concentration of ∼14 mg/mL at -80 ºC until use. Yields were 95 mg purified AS protein/L growth medium for the uniform [^13^C, ^15^N] labeled monomer.

### Preparation of Insoluble fraction seeds from LBD, MSA and control postmortem tissue

The protocol to sequentially extract human postmortem brain tissue was adapted from Appel-Cresswell et al^2^. Briefly, gray matter dissected from tissue was sequentially homogenized in four buffers (3 ml/g wet weight of tissue) using Kimble Chase Konte™ dounce tissue grinders (KT885300-0002). In the first step, 300mg of dissected grey matter tissue was homogenized using 20 strokes of Pestle A in High Salt (HS) buffer (50 mM Tris-HCl pH 7.5, 750 mM NaCl, 5 mM EDTA plus Sigma P2714 Protease Inhibitor (PI) cocktail). The homogenate was centrifuged at 100,000 ×g for 20 min at 4 °C and the pellet was homogenized in the next buffer using 20 strokes of Pestle B. Extractions using Pestle B were performed in HS buffer with 1% Triton X-100 with PI, then HS buffer with 1% Triton X-100 and 1 M sucrose, and with 50 mM Tris-HCl, pH 7.4 buffer. In the final centrifugation, the resulting pellet was resuspended in 50 mM Tris-HCl, pH 7.4 buffer (3 ml/g wet weight of tissue). The aliquots of insoluble fraction were stored at -80 °C until use. Similar extraction protocol was followed for LBD, MSA and control cases.

### Amplification of isotopically labelled LBD fibrils from LBD insoluble fraction seeds

We amplified LBD-fibrils from gray matter dissected from the caudate region. We incubated insoluble fraction seeds with an Asyn monomer preparation containing isotopically labeled Asyn monomer supplemented with control fraction. The control fraction preparation was derived from E.Coli transformed with an empty expression vector, and was purified with the same protocol as the natural abundance Asyn monomer (JBC, 2017 V292, Pg9034). Asyn monomer and control fraction was filtered through a 50k MWCO Amicon Ultra centrifugation filter (Millipore, UFC805204) before use, to remove any preformed aggregates.

Insoluble fraction (10 µL) containing 3.3 µg wet wt. of tissue was bought to a final volume of 30 µL by addition of 20 mM Tris-HCl, pH 8.0 plus 100 mM NaCl buffer (fibril buffer) in a 1.7mL microcentrifuge tube. The insoluble fraction was sonicated for 2 min at amplitude 50 in a bath sonicator (Qsonica model Q700) with a cup horn (5.5 inch) attachment at 4 °C. To the sonicated seeds, 1.5 µL of 2 % Triton X-100 was added. To this mixture, 50k Amicon ultra filtered isotopically labelled Asyn monomer was added to a final concentration of 2 mg/mL in a final volume of 100 µL. This mixture underwent quiescent incubation at 37 °C for 3 days, completing the 1^st^ round of sonication plus incubation. After the first round, the mixture was sonicated at 1 min at amplitude 50, and then an additional 300 µL of 2 mg/mL Asyn monomer was added. The mixture underwent quiescent incubation at 37 °C for 2 days (2^nd^ round). Then, sonication for 1 min at amplitude 50 and quiescent incubation for 2 days was repeated for the third round, followed by sonication for 1 min at amplitude 50 and quiescent incubation for 3 days for the 4^th^ round. At the end of 4^th^ round, LBD-amplified fibrils were stored at 4 °C until use.

Further expansion of the LBD-amplified fibrils was performed by centrifuging 60 μL of 4^th^ round LBD-amplified fibrils at 21,000 xg for 15 min at 4 °C. The pellet was resuspended in 100 µL of fibril buffer and sonicated for 1 min at amplified 50. To this mixture, Asyn monomer was added to a final concentration of 2 mg/mL in a final volume of 400 µL in fibril buffer. This mix was quiescently incubated at 37 °C for 2 days (5^th^ round). At the end of 5^th^ round, samples were centrifuged at 21,000 xg for 15 min at 4 °C and the top 300 µL of spent Asyn monomer was moved to a separate tube. The pellet was resuspended by trituration and sonicated for 1 min at amplified 50. After sonication, the previously removed 300 µL of 5^th^ round monomer was added back. Next, an additional 2.5 mg/mL of Asyn monomer was added to bring the total volume to 800 µL. This mixture was incubated at 37 °C for 2 days to complete 6^th^ round of incubation. The increased monomer concentration (2.5 mg/mL instead of 2 mg/mL) was calculated based on the average decrease in free Asyn monomer due to its incorporation into amplified fibrils. The 6^th^ round fibrils were stored at 4 °C until use.

### Solid-state NMR spectroscopy

Magic-angle spinning (MAS) SSNMR experiments were performed at magnetic field of 11.7 T (500 MHz ^1^H frequency) or 17.6 T (750 MHz ^1^H frequency) using Agilent Technologies VNMRS spectrometers. Spinning was controlled with a Varian MAS controller to 11,111 ± 30 Hz or 22,222 ± 15 Hz (11.7 T) and 16,667 ± 15 Hz or 33,333 ± 30 Hz (17.6 T), with two minor exceptions indicated in Table 1. All experiments were done with a variable-temperature (VT) airflow setting of 0 ºC, primarily to keep samples cool from RF and MAS heating, without freezing out molecular motions. The 11.7 T magnet was equipped with a 1.6 mm HCDN T3 probe (Varian), with pulse widths of about 1.8 µs for ^1^H and ^13^C, and 3.2 µs for ^15^N. The 17.6 T magnet was equipped with a HXYZ T3 probe (Varian) tuned to HCN triple resonance mode with pulse widths of about 1.9 µs for ^1^H, 2.6 µs for ^13^C, and 3.0 µs for ^15^N. All experiments utilized ^1^H-^13^C or ^1^H-^15^N tangent ramped CP (Metz et al. 1994) and ∼100 kHz SPINAL-64 decoupling during evolution and acquisition periods (Comellas et al. 2011, Fung et al. 2000). Where applicable, SPECIFIC CP was used for ^15^N-^13^Cα and ^15^N-^13^C’ transfers (Baldus et al. 1998), ^13^C-^13^C homonuclear mixing was performed using DARR (Takegoshi et al. 2001). Chemical shifts were externally referenced to the downfield peak of adamantane at 40.48 ppm (Morcombe and Zilm 2003). NUS schedules using biased exponential sampling were prepared using the nus-tool application in NMRbox (Maciejewski et al. 2017). Data conversion and processing was done with NMRPipe (Delaglio et al. 1995). NUS data was first expanded with the nusExpand.tcl script in NMRPipe, converted, and processed using the built-in SMILE reconstruction function (Ying et al. 2017). Peak picking and chemical shift assignments were performed using NMRFAM-Sparky (Lee et al. 2015).

**Table 1:** Table 4.2 ^13^C, and ^15^N, chemical shift assignments for LBD derived α-synuclein fibrils

| Residue | $^{15}\text{N}$ | $^{13}\text{C}'$ | $^{13}\text{CA}$ | $^{13}\text{CB}$ | $^{13}\text{CG}$ | $^{13}\text{CD}$ | $^{13}\text{CE}$ | $^{13}\text{CZ}$ | $^{15}\text{ND}/^{15}\text{NE}/^{15}\text{NZ}$ |
| --- | --- | --- | --- | --- | --- | --- | --- | --- | --- |
| --- | --- | --- | --- | --- | --- | --- | --- | --- | --- |
| K34 | 125.5 | 174.5 | 56.1 | - | - | - | - | - | - |
| E35 | 121.9 | 176.3 | 53.4 | 35.7 | 35.6 | 183.4 |  |  |  |
| G36 | 117.5 | 173.6 | 48.4 |  |  |  |  |  |  |
| V37 | 118.6 | 172.4 | 60.0 | 35.7 | 21.9/24.5 |  |  |  |  |
| L38 | 127.8 | 173.8 | 52.6 | 47.3 | 27.5 | 25.3/29.0 |  |  |  |
| Y39 | 131.3 | 173.9 | 56.3 | 43.4 | 128.0 | 133.0 | 117.5 | 157.3 |  |
| V40 | 127.1 | 173.8 | 60.0 | 34.4 | 19.9/22.1 |  |  |  |  |
| G41 | 111.6 | 170.2 | 44.7 |  |  |  |  |  |  |
| S42 | 116.9 | 175.4 | 56.5 | 68.7 |  |  |  |  |  |
| K43 | 123.4 | 176.6 | 56.6 | - | - | - | - | - | - |
| --- | --- | --- | --- | --- | --- | --- | --- | --- | --- |
| V49 | - | - | - | - |  |  |  |  |  |
| H50 | - | 175.8 | 51.7 | 31.5 | 125.1 | 129.4 | 140.3 |  | -/- |
| --- | --- | --- | --- | --- | --- | --- | --- | --- | --- |
| E61 | - | 173.1 | - | - | - | - |  |  |  |
| Q62 | 122.6 | 174.3 | 54.5 | - | - | - |  |  | - |
| V63 | 124.6 | 173.9 | 61.2 | 35.5 | 21.1/- |  |  |  |  |
| T64 | 124.8 | 172.3 | 60.8 | 68.5 | 23.4/23.4 |  |  |  |  |
| N65 | 126.5 | 173.0 | 51.9 | 42.0 | 174.4 |  |  |  | 113.9 |
| V66 | 125.0 | 177.0 | 59.9 | 34.4 | 20.9/23.0 |  |  |  |  |
| G67 | 112.6 | 171.9 | 47.4 |  |  |  |  |  |  |
| G68 | 100.7 | 172.1 | 44.4 |  |  |  |  |  |  |
| A69 | 121.8 | 175.0 | 50.0 | 22.1 |  |  |  |  |  |
| V70 | 124.5 | 175.8 | 61.7 | 32.7 | 19.7/21.2 |  |  |  |  |
| V71 | 131.0 | 173.9 | 61.3 | 34.7 | 19.7/22.1 |  |  |  |  |
| T72 | 124.7 | 174.6 | 60.1 | 71.4 | 22.5 |  |  |  |  |
| G73 | 117.8 | 172.1 | 49.0 |  |  |  |  |  |  |
| V74 | 117.6 | 173.8 | 59.4 | 37.4 | 20.3/23.3 |  |  |  |  |
| T75 | 123.4 | 171.8 | 61.7 | 68.8 | 22.9 |  |  |  |  |
| A76* | 131.2 | 174.2 | 49.7 | 21.7 |  |  |  |  |  |
| V77* | 122.4 | 172.2 | 60.6 | 36.7 | 20.6/25.1 |  |  |  |  |
| A78 | 130.3 | 175.7 | 50.7 | 23.1 |  |  |  |  |  |
| Q79 | 122.6 | 174.1 | 54.6 | 37.6 | 34.7 | 179.3 |  |  | 112.4 |
| K80 | 126.5 | 173.9 | 54.7 | - | - | - | - | - | - |
| T81 | 124.7 | 172.8 | 61.9 | 69.9 | 21.1 |  |  |  |  |
| V82 | 127.0 | 175.4 | 60.1 | - | -/- |  |  |  |  |
| E83 | 118.3 | 175.9 | 55.3 |  |  |  |  |  |  |
| G84 | 124.4 | 175.0 | 49.8 |  |  |  |  |  |  |
| A85 | 124.7 | 174.5 | 49.9 | - |  |  |  |  |  |
| G86 | 110.3 | 172.5 | 44.2 |  |  |  |  |  |  |
| S87* | 119.7 | 174.4 | 56.4 | 67.4 |  |  |  |  |  |
| I88* | 128.2 | 173.2 | 60.1 | 44.4 | 27.8/17.6 | 16.9 |  |  |  |
| A89 | 129.1 | 175.2 | 50.3 | 22.5 |  |  |  |  |  |
| A90 | 126.3 | 174.4 | 49.7 | 22.2 |  |  |  |  |  |
| A91 | 123.7 | 175.4 | 50.3 | 23.2 |  |  |  |  |  |
| T92 | 117.3 | 173.3 | 61.2 | 70.8 | 20.5 |  |  |  |  |
| G93 | 116.8 | 170.3 | 44.9 |  |  |  |  |  |  |
| F94 | 129.0 | 173.1 | 56.9 | 43.5 | 137.4 | 131.2 | 129.3 | 140.3 |  |
| V95 | 128.5 | 173.0 | 60.9 | 35.6 | -/- |  |  |  |  |
| K96 | 129.0 | 175.1 | 54.5 | - | - | - | - | - | - |

(\*) Two sets of shifts observed for at least one atom in the residue. Primary shift is reported here.

Assignments and data deposition

### Chemical Shift Assignments and Extent of Assignments

Chemical shift assignments were performed for the LBD Asyn fibrils amplified using uniform [^13^C, ^15^N] labeling (uCN). Resonance assignments were determined using 2D ^13^C-^13^C, 2D ^15^N-^13^Cα, 2D ^15^N-^13^C’, 3D ^15^N-^13^Cα-^13^CX, 3D ^15^N-^13^C’-^13^CX, 3D ^15^N-^13^C’-^13^Cα and 3D ^13^Cα-^15^N-^13^C’ data sets following standard procedures (Comellas and Rienstra, 2013; Higman, 2018). The complete list of data sets used for performing assignments using ^13^C-detection is presented in Supplementary Table 1. The 2D ^13^C-^13^C spectrum serves as a conformational fingerprint of the fibril and provides some initial insights into the structure Figure 1A. Globally, the resolved peaks display line widths, including scalar couplings, of < 0.4 ppm in the direct ^13^C dimension, similar to those observed for *in vitro* Asyn fibrils and indicative of a highly ordered core (Barclay et al., 2018; Comellas et al., 2011a). Particular residue types including Thr, Val, and Gly, as well as select spin systems including L38, N65, Q79, I88, and F94 are particularly well resolved in the 2D. The Lys and Glu regions, which account for 20% of the primary sequence between residues 30 and 100, are broad and poorly resolved as with the *in vitro* form, indicating disorder or partial mobility for the majority of these residues. In contrast to the *in vitro* form, the Ala regions are poorly resolved in the 2D ^13^C-^13^C, but the signals are of comparable intensity to the resolved regions, indicating that the alanines are highly ordered but of similar structure throughout the core. Interestingly, S87 displays two sets of peaks that combine to approximately half of the intensity of the strong peaks, while I88 shows broadening for the peaks arising from backbone correlations, suggesting at least two conformations for this region. In the 3D spectra, a few additional residues, namely A76 and V77, also show two sets of peaks but for the side chain atoms. In addition, the Asyn sequence contains 10 Thr residues all located between residues 22 and 92, but at least 11 appear in the 2D ^13^C-^13^C spectrum. The 2D ^15^N-^13^C’ (Figure 1A) and ^15^N-^13^Cα (Figure 1B) spectra also serve as structural fingerprints for the fibril, with a focus on the backbone atoms. Critically, these highlight the benefit of adding a ^15^N dimension to disambiguate shifts, particularly for the Ala and Gly regions, which comprise 30% of the primary sequence between residues 30 and 100, as well as for key core residues like V71, V74, T75 and V77. Overall, there appears to be a predominant defined conformation that likely displays some localized heterogeneity. Figure 2 demonstrates representative strips corresponding to assignments from 3D ^15^N-^13^Cα-^13^CX, 3D ^15^N-^13^C’-^13^CX, and 3D ^13^Cα-^15^N-^13^C’ from T72 to A76.

**Figure 1:**
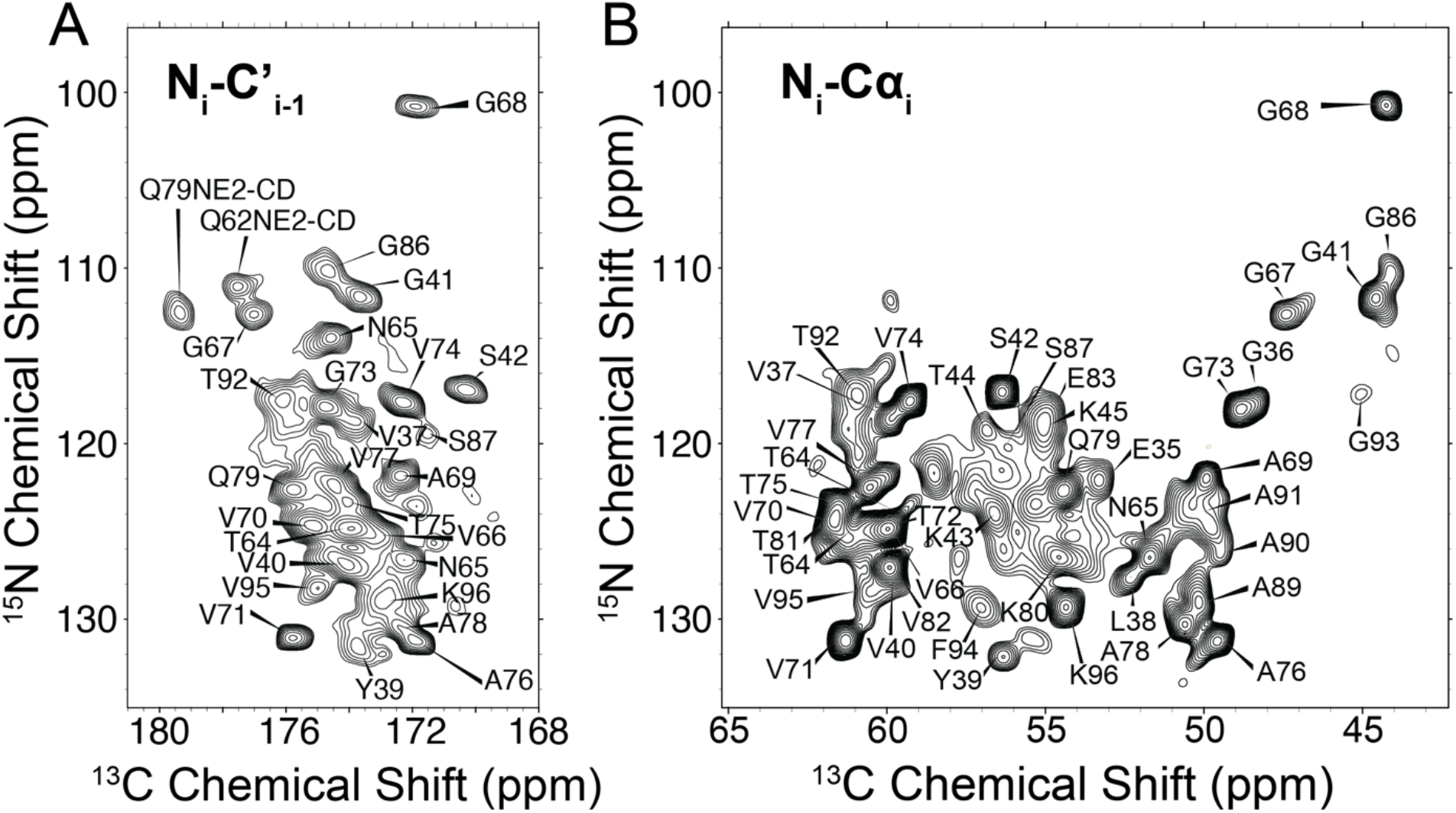
Backbone chemical shift assignments demarked on a ^15^N-^13^C’ (A) and a ^15^N-^13^Cα (B) spectra

**Figure 2:**
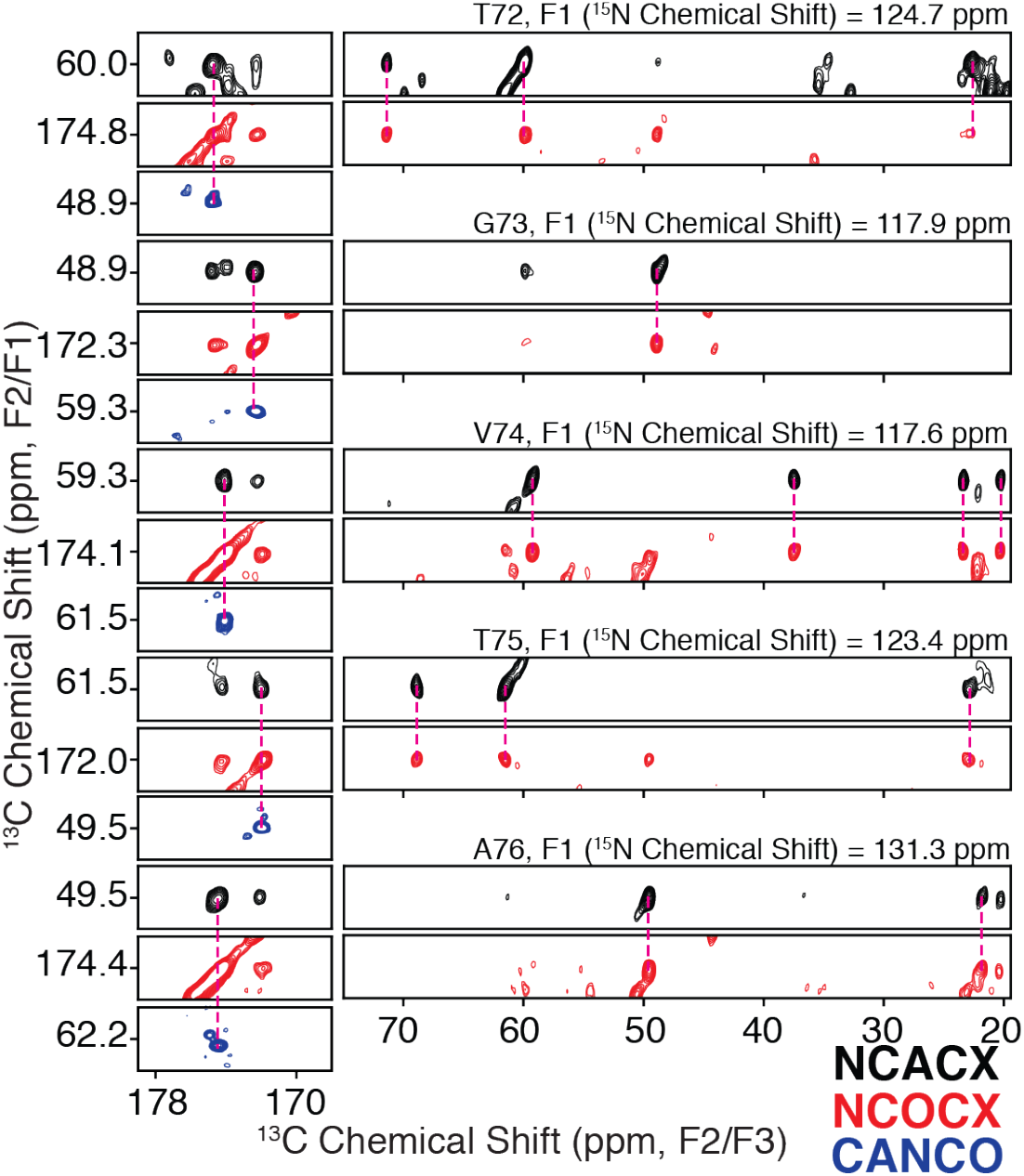
Example backbone assignment strip from T72 to A76 demonstrating connectivity and sidechain assignments from a 3D ^15^N-^13^Cα-^13^CX correlation (black), a 3D ^15^N-^13^C’-^13^CX correlation(red), and a 3D ^13^Cα-^15^N-^13^C’correlation (blue).

## Data availability

Resonance assignments are available through the Biological Magnetic Resonance Data Bank (http://bmrb.io), Accession Number 51678.

